# Thickening of the cell wall increases the resistance of *S. cerevisiae* to commercial formulations of glyphosate

**DOI:** 10.1101/760694

**Authors:** Apoorva Ravishankar, Amaury Pupo, Jennifer E.G. Gallagher

## Abstract

The use of glyphosate-based herbicides is widespread and despite its extensive use, its effects are yet to be deciphered completely. The additives in commercial formulations of glyphosate, though labeled as inert when used individually, have adverse effects when used in combination with other additives and the active ingredient. As a species, *Saccharomyces cerevisiae* has a wide range of resistance to glyphosate-based herbicides. To investigate the underlying genetic differences between sensitive and resistant strains, global changes in gene expression were measured when yeast were exposed to a commercial formulation of glyphosate (CFG). Changes in gene expression involved in numerous pathways such as DNA replication, MAPK signaling, meiosis, and cell wall synthesis. Because so many diverse pathways were affected, these strains were then subjected to in-lab-evolutions (ILE) to select mutations that confer increased resistance. Common fragile sites were found to play a role in adaptation mechanisms used by cells to attain resistance with long-term exposure to CFG. The cell wall structure acts as a protective barrier in alleviating the stress caused by exposure to CFG. The thicker the cell wall, the more resistant the cell is against CFG. Hence, a detailed study of the changes occurring at the genome and transcriptome level is essential to better understand the possible effects of CFG on the cell as a whole.

**Author Summary:** We are exposed to various chemicals in the environment on a daily basis. Some of these chemicals are herbicides that come in direct contact with the food we consume. This makes the thorough investigation of these chemicals crucial. Some of the most commonly used herbicides around the world are glyphosate-based. Their mode of action effects a biosynthetic pathway that is absent in mammals and insects and so it is deemed safe for consumption. However, there are many additives to these herbicides that increase its effects. Thorough testing of these commercially available herbicides is essential to decipher all the potentially adverse effects that it could have on a cell. *Saccharomyces cerevisiae* has a wide range of genetic diversity, making it is suitable to test different chemicals and identify any harmful effects. In this study, we exposed yeast cells to some glyphosate-based herbicides available in the market, to understand what effects it could have on a cell. We found that the additives in the herbicides have an effect on the cell wall and the mode of entry of glyphosate into the cell.

## Introduction

Glyphosate-based herbicides are one of the most commonly used broad-spectrum herbicides around the world [1] because of its low toxicity to mammals, high efficacy and affordability in comparison to other herbicides [2,3]. Glyphosate acts by inhibiting the aromatic amino acid biosynthesis produced by the shikimate pathway. It targets the 5-enolpyruvylshikimate-3-phosphate synthase (EPSPS) enzyme that plays a crucial role in the conversion of phosphoenolpyruvate (PEP) and 3-phosphoshikimate to phosphate and 5-enolpyruvylshikimate-3-phosphate (Amrhein, et al. 1980; Powles & Preston, 2006). Glyphosate binds to EPSPS in plants, in turn inhibiting the shikimate pathway and the production of tryptophan (W), tyrosine (Y), phenylalanine (F), parabenozic acid and Coenzyme Q10 (Haderlie et al. 1977).

Organisms undergo different modes of adaptation to attain resistance to an environmental stressor. In the case of glyphosate-based herbicides, the routes to attaining resistance are classified into two categories: 1. Target site associated resistance, and 2. Non-target site resistance (Yuan, et al. 2007). Target site associated resistance is comprised of resistance mechanisms involving changes in the EPSPS gene and the shikimate pathway. Typically, either the glyphosate binding site in EPSPS is mutated or EPSPS is overexpressed. Non-target site resistance is attained due to changes outside the shikimate pathway. One of the most commonly studied form of non-target site resistance is vacuole related changes (Hereward et al. 2018; Moretti et al. 2017). Recent studies showed that genes involved in glyphosate transport are one of the non-target resistance mechanisms (Rong-Mullins et al. 2017). Some yeast strains encode a CGF resistant pleiotropic drug response protein, Pdr5 (Rong-Mullins et al. 2017). The Pdr5 protein is involved in the transport of glyphosate and CGF additives out of the cell. Another non-target resistance mechanism comes from the protein Dip5, a glutamic and aspartic acid permease and its deletion leads to glyphosate tolerance (Rong-Mullins et al. 2017).

There are hundreds of commercial formulations of glyphosate (CFG) available in the market. The most challenging aspect of studying the effects of these CFG is that the label lists the concentration of glyphosate present in the mixture but does not provide details regarding the additives added or the concentrations they are added in. It is crucial to monitor and study these CFG as a whole, along with the additivess, as they alter the effectiveness of the principal component and other factors such as its extent of biodegradability (Mesnage et al. 2013). Some of the identified additives are polyoxyethylamines (POE), quaternary ammonium compounds and some heavy metals (Defarge et al. 2018; Nicolas Defarge et al., 2016; Nørskov et al. 2019). As the contributions of these additives to the toxicity of the herbicide are now being studied in more detail [15]. How these additives inflict changes on the genome and transcriptome of organisms would vary depending of the specific CFG. A diverse group of stressors triggers the environmental stress response (ESR) which is seen in the change of global yet similar pattern of gene expression. The changes in transcript levels of genes observed in the state of ESR are involved in various cellular processes such as energy generation and storage, defense against reactive oxygen species, protein folding, DNA repair, etc. The protection of these pathways are vital to the proper functioning of the cell [16].

The yeast cell wall is 15-30% of the dry weight of the cell [17]. The cell wall is mainly comprised of an inner layer made of polysaccharides and an outer scaffold made of mannoproteins (Klis, et al. 2002; Orlean, 2012; Stewart & Stewart, 2018). The mannoproteins, β-1,3 glucans, and β-1,6 glucans form the major components of the cell wall, with chitin forming non-covalent bonds with some glucans as a minor component. The cell wall is highly dynamic in nature and it can adapt to various physiological and morphological conditions [21]. The function of the cell wall is to stabilize internal osmotic pressure, protect the cell against mechanical injury and chemical stress, maintain the cell shape, and provide a scaffold for glycoproteins [20]. The outer protein layer of the cell wall contains at least 70 proteins, which may vary depending on the environment and the type of stressors the cells are exposed to (De Groot et al. 2005; Zlotnik et al. 1984). Proteins containing a large amount of N- and O-linked mannoses are called mannoproteins, many of which are found on the outermost layer of the cell wall [23]. Some of these structural cell wall proteins are rich in serine and threonine and undergo a post-translational addition of a glycosyl phosphatidylinositol (GPI) anchor (Nuoffer et al. 1991). Sed1 is one of many GPI-anchored mannoproteins in the cell wall-bound to a glucan. Stationary phase cells show an increase in Sed1 levels (Shimoi et al. 1998). Cells in stationary phase tend to be much more resistant to various environmental stressors (Werner-Washburne et al. 1993) and induction of these proteins is usually observed on exposure to stress [25].

*Sacchromyces cerevisiae* has a wide range of genetic variation within the species. Some of this variation pertains to the origin of these strains, in terms of their prior exposure to different stressors. In this study, we used the genetic variation in yeast to assess the effects of CFG on the transcriptome of different strains of yeast. We also analyzed various modes of attaining resistance to CFG that lie outside the shikimate pathway. CFG caused a pleiotropic response on cells exposed to it. They affected numerous pathways involved in various vital processes for the cell’s survival such as meiosis, DNA replication, cell wall proteins, MAPK and HOG signaling pathways, etc. The high osmotic response (HOG pathway) aids in the survival of the cell in case of cell wall stress (García et al. 2009). Cell wall structure plays an important role in resistance to CFG. Some cell wall proteins such as Sed1, when expressed in high numbers contributed to the thickness of the cell wall and aided in tolerance to CFG. Another interesting aspect was the contribution of Ty elements to the adaptation of tolerance mechanisms against CFG. Numerous genes that lie between Ty elements underwent duplication during the process of developing adaptation mechanisms on CFG exposure. Many of these changes could have a cumulative response in the cells developing resistance against CFG.

## Results

### Variation in growth in the presence of glyphosate-based commercial formulations

Four genetically diverse haploid strains with well-annotated genomes (S1 Table [28]) were chosen to test for differences in growth when exposed to different commercial formulations of glyphosate (CFG). RM11 is a vineyard strain [29,30] and AWRI1631 is descended from commonly used strain for commercial wine manufacturing [31]. YJM789 is derived from a clinical isolate, isolated from an immunocompromised AIDS patient (McCusker et al. 1994; Tawfik et al. 1989). The last strain included was a commonly used laboratory strain with an S288c background (GSY147) (Engel et al., 2014a; R K Mortimer & Johnston, 1986; Wenger et al. 2010). These strains were exposed to commercial formulations of glyphosate-based herbicides on solid media, with pure glyphosate as a control. We exploited this variation to decipher the effects of the different commercially available formulations of glyphosate-based herbicides. There are a large number of formulations available, many of which contain a mixture of multiple active ingredients such as pelargonic acid, diquat, imazapic, and other chemicals. The ones selected for this study were chosen because they were labeled as containing glyphosate as the sole active ingredient in the herbicide, hence the variation observed is the clear effect of the putatively inert ingredients such as addivates.

When grown with CFG on rich media (YPD), AWRI1631 and RM11 continued to grow in the presence of 1% glyphosate in all the formulations (Fig 1A). Whereas YJM789 and S288c displayed a growth defect, and S288c was extremely sensitive to only Weed Pro (WP) as compared to the other three formulations. On minimal media (YM), where cells synthesize their own amino acids from the nitrogen-base provided, ammonium sulfate, in this case, there was greater variation between the formulations, especially in the case of WP where all the strains were sensitive, but more or less the pattern was consistent among the other formulations, with S288c showing maximum sensitivity and AWRI1631 showing resistance to a larger extent (Fig 1B). As glyphosate targets, the aromatic amino acid pathway, supplementing the minimal media with tryptophan, tyrosine, and phenylalanine (WYF) should bypass the inhibition of the shikimate pathway and restore growth. However, rescue with WYF and the CFG was variable, and growth improved to a great extent; although the cells did not fully recover (Fig 1C). AWRI1631 and RM11 growth inhibition was less compared to YJM789 and S288c. S288c was the most affected by all the formulations. All the strains were sensitive to WP and supplementing WYF did not show considerable rescue in this formulation. This indicates that there are other factors influencing the cell growth other than the active ingredient, and these vary from one formulation to the other. Dip5 is an amino acid permease that allows entry of glyphosate into the cell [10]. It is downregulated in the presence of excess aspartic acid (Hatakeyama et al. 2010) and adding back aspartic acid did rescue the cells significantly, but not entirely in all the formulations, especially in case of S288c (Fig 1D). The addition of excess aspartic acid to cells treated with pure glyphosate rescued the growth inhibition. In the case of CFG, the alleviation of growth inhibition was not much with the addition of aspartic acid. YJM789 was an exception to this, as there was a significant rescue on treatment with Credit 41 (Cr41) when supplemented with aspartic acid. The growth inhibition resulting from exposure to WP was alleviated to a greater extent on the addition of aspartic acid in comparison to WYF, but not completely. In comparison, when yeast were grown on pure glyphosate (PG) the variation in growth was similar to that of Cr41, Compare and Save (CAS), and Super Concentrated (SC) but to a lesser extent, which supports the hypothesis that the additives do contribute to growth inhibition. YJM789 was the exception as it was the most sensitive to the pure glyphosate in all conditions. To consider a formulation that is representative of a phenotype similar to most commercial formulations, Cr41 was used for all further experiments.

**Fig 1:**
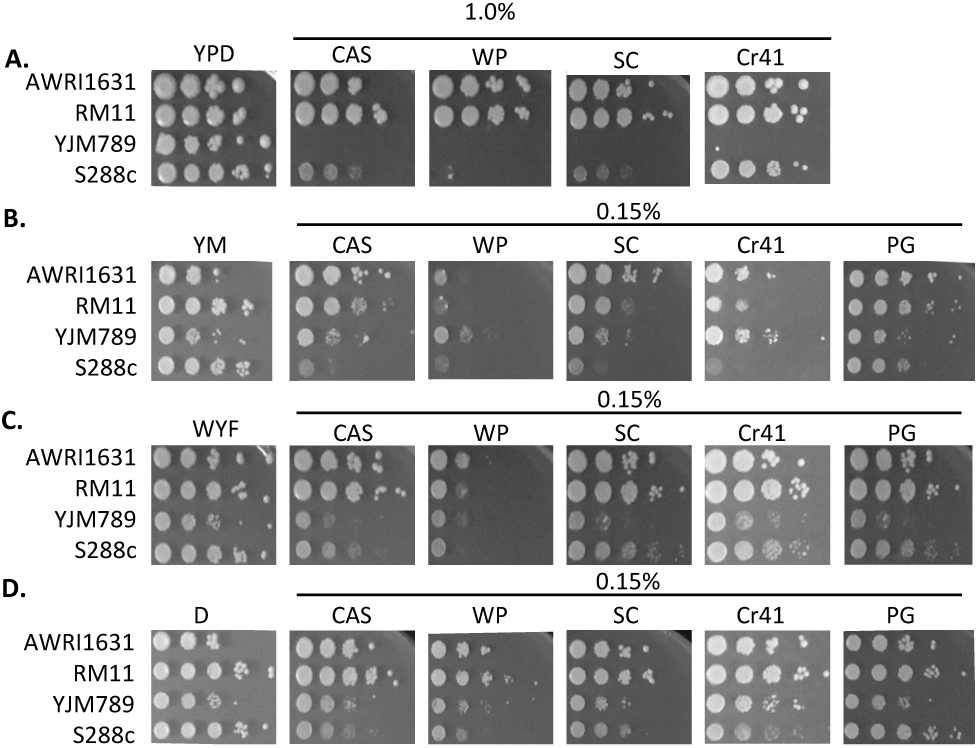
Contribution of genetic variation of different yeast strains to glyphosate resistance. Serial dilutions of haploid AWRI1631, RM11, YJM789, and S288c were grown on the following media **A**. YPD with 1.0% of different formulations of glyphosate-based herbicides. **B**. YM with 0.15% of different formulations of glyphosate-based herbicide. **C**. YM+ aromatic amino acids (WYF) with 0.15% of different formulations of glyphosate-based herbicides and **D**. YM+ aspartic acid (D) with 0.15% of different formulations of glyphosate-based herbicides. Compare and Save (CAS); WeedPro (WP); Super Concentrated (SC); Credit 41 (Cr41) and pure glyphosate (PG).

### A polygenic trait that has a strain and condition-based pattern in response to Cr41

There are tens of thousands of SNPs between these four strains. To explore how genomes change to permit adaptation to high levels of Cr41, the yeast were serially passaged in media containing Cr41 by performing In-Lab-Evolutions (ILEs). Three biological replicates were passaged in YM, YM supplemented with WYF and YPD media, with 0.25% glyphosate from the Cr41 in minimal media and 1% in rich media respectively (Fig 2A). The strains were diluted through six passages by transferring 1% culture to fresh media. Once the resistant populations were identified, single colonies were isolated. These strains were then released from the selective pressure for two passages and then the Cr41 resistance was confirmed (Fig 2B-E). This step was performed in order to ensure that the resistance wasn’t entirely due to epigenetic mechanisms that cannot be detected through whole-genome sequencing. The resistant cells sequenced for each strain along with the condition are listed in S2 Table. The ILEs selected a large number of SNPs as well as duplicated regions. The synonymous mutations were filtered out and only the genes containing non-synonymous mutations were taken into consideration. A total of 148 genes (S3 Table) accumulated at least one non-synonymous SNP within the coding region among all the sequenced strains treated with Cr41. The genes that accumulated more than one SNP and/or indel in any of the samples from different strains were prioritized and focused on for this study. Dip5 in part transports glyphosate into the cell as shown in our previous study [10], and contained SNPs in three of the sequenced samples (Fig S1).

**Figure 2:**
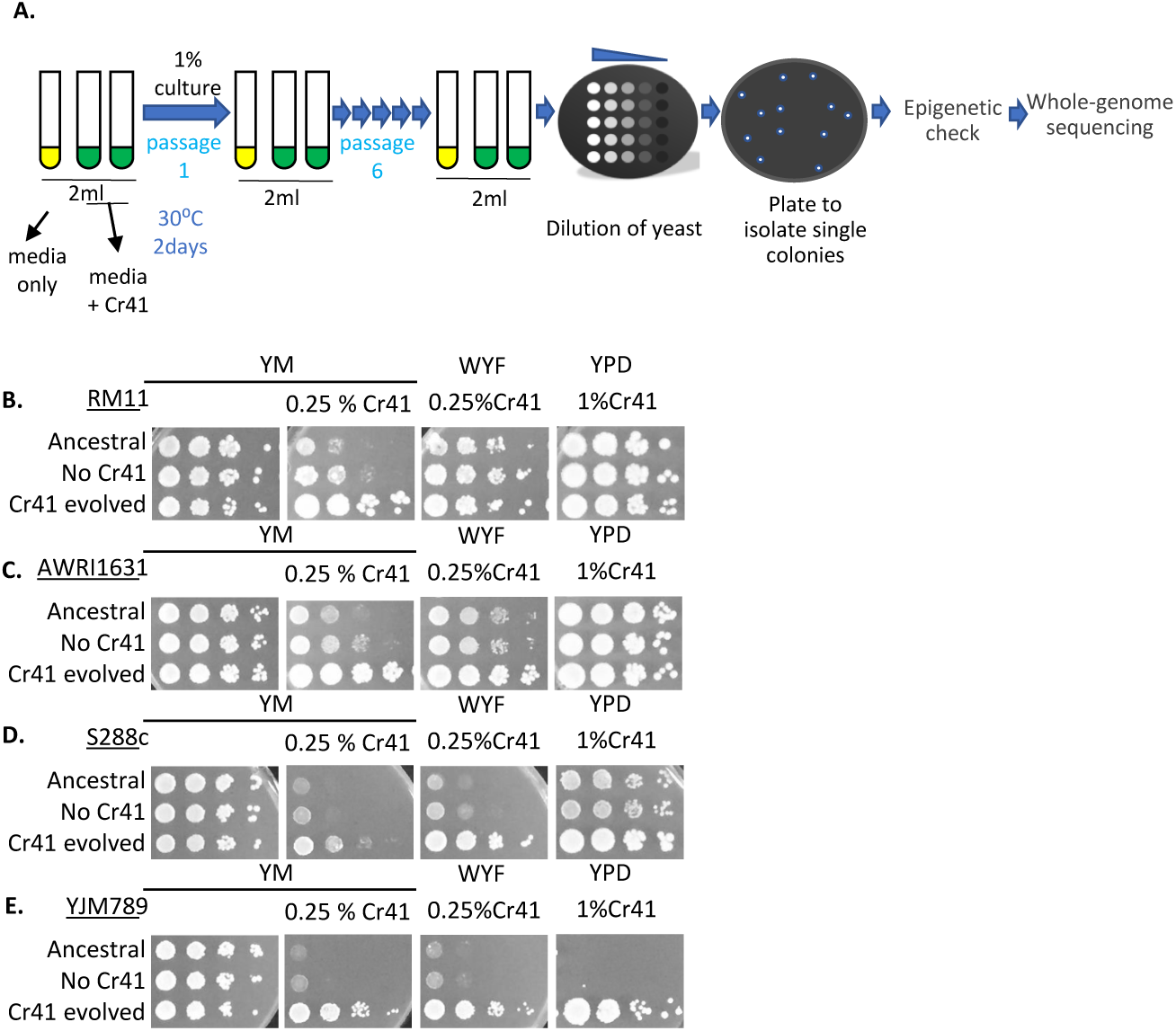
Selection of strains resistant to commercial glyphosate formula, Credit41 by In-lab evolutions (ILEs). **A**. Outline of In-lab evolutions (ILEs) methodology: The cells are grown in media with and without Credit41 for 2 days at 30°C on the shaker. 1% of the culture was transferred to fresh media for 6 passages. Serial dilutions of the cultures of the sixth passage were plated on solid media. The resistant lines were plated on solid media to isolate single colonies. The genomic DNA was extracted and submitted for Illumina sequencing. **B**. Serial dilutions of haploid RM11 strain evolved without Cr41 and strain evolved with Cr41, on media with Cr41. **C**. Serial dilutions of haploid AWRI1631 strain evolved without Cr41 and strain evolved with Cr41 on media with Cr41. **D**. Serial dilutions of haploid S288c, strain evolved without Cr41 and strain evolved with Cr41 on media with Cr41. **E**. Serial dilutions of haploid YJM789, str ain evolved without Cr41 and strain evolved with Cr41 on media with Cr41.

Principal Component Analysis (PCA plots) were plotted for each sample taking all the SNPs into consideration in order to identify the major sources of variation. This was done to ensure that the main sources of variation corresponds to the biological conditions and was not a mere effect of experimental bias. An R-package, pcadapt (Luu et al. 2017) visualized the patterns that naturally exist between the strains. Hence, resistant strains clustered together and so did the sensitive strains when the data from all the conditions were pooled together (Fig 3A). The underlying background genetic differences dominated the genome comparisons. Further analysis of the strain-specific PCA plots showed that the clustering of the controls vs. the Cr41 treated samples in S288c was based on PC2 (Fig 3B), and PC1 did the same in the case of RM11 (Fig 3C).

**Figure 3:**
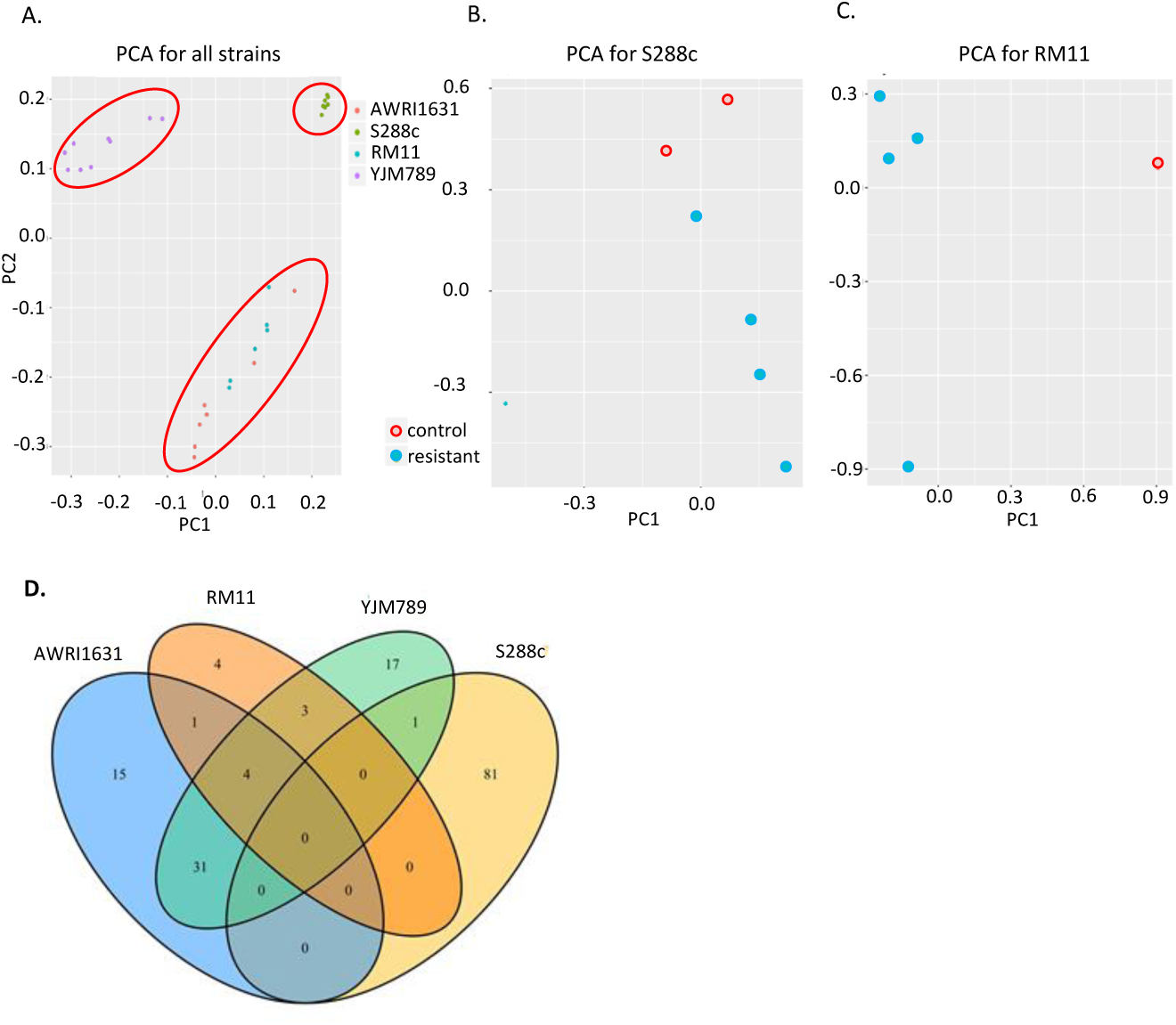
Whole-genome analysis of In-lab evolutions (ILEs). **A**. PCA of SNPs of various samples from all 4 strains, AWRI1631, S288c, RM11, and YJM789. **B**. PCA of SNPs identified among the S288c ILE samples **C**. PCA of SNPs identified among the RM11 ILE samples. **D**. Venn-diagram representing all the affected genes i.e., those that contained CNV as well as those that accumulated SNPs, in each strain.

Sections of the genome containing as few as two to as many as a hundred genes increased in copy number (CN) as shown by copy number variation analysis (CNV). All the strains in this study were haploid and having a CN of less than one can only occur at nonessential genes. In this analysis, any genes that were partially duplicated were excluded under the assumption that the change in copy number is associated with synteny or the other genes present in their immediate surroundings (S4 Table). Any gene that underwent a CNV in a treated cell but also had an increase in CN in any control sample, was filtered out. This made it apparent that there is an underlying connection between the strain and the condition (i.e. YM, YM with WYF or YPD) it was evolved under. As in the case of YJM789, CNVs occurred only in cells that were evolved in minimal media with WYF. S288c had the most genes (81 genes) that underwent CNV while RM11 did not have any, which was one of the most CFG resistant strain. Some of the S288c genes that underwent CNV were found in all the conditions, but many of which were found in cells evolved in minimal media supplemented with WYF. Though these genes underwent duplication in a specific condition (i.e., WYF), they did not belong to a single pathway. Their functionality ranged from mitochondrial maintenance as in the case of *MRX14, MRLP1* to biosynthesis of secondary metabolites such as *INO2*. This led to the hypothesis that the route used to attain resistance was dependent on the media type. Hence, if a certain strain was evolved in YM, it may not be resistant to the CFG in YPD. To confirm this hypothesis, semi-quantitative growth assays of the evolved strains was carried out in different combinations of media conditions. The strains evolved in rich media did not confer cross-resistance in minimal media. However, those evolved in minimal media and WYF concur cross-resistance (Fig S2).

Many regions that underwent CNV were flanked by fragile sites, mainly consisting of transposable elements [39] and one long regulatory ncRNA, *ICR1* (Table 1). Fragile sites are defined as regions of the genome that make it difficult for the cell to undergo replication and sometimes result in chromosome breakage. Studies in other organisms such as *C*. *elegans* and human cells have shown evidence of DNA damage in these sites [15,40]. Past studies lack conclusive evidence to declare that Ty elements are involved in fragile site rearrangements resulting in CNV of certain regions. However, there is evidence to prove that these elements are involved in processes such as translocations, deletions, etc. (Dunham et al. 2002; Roeder & Fink, 1980). Most of the regions undergoing an increase in copy number and being flanked by Ty elements were found in S288c cells, evolved in the presence of Cr41. All the genes that contained non-synonymous SNPs or underwent CNV (S4 Table) were referred to as affected genes because they were affected by the Cr41 treatment (Fig 3D). The regions that underwent CNV were large sections containing multiple genes from different pathways such as MAP kinase/ Hog1 pathway, mitochondrial genes, DNA damage repair pathways, spindle formation, metal transporters, cell wall, and cell membrane.

**Table 1.**
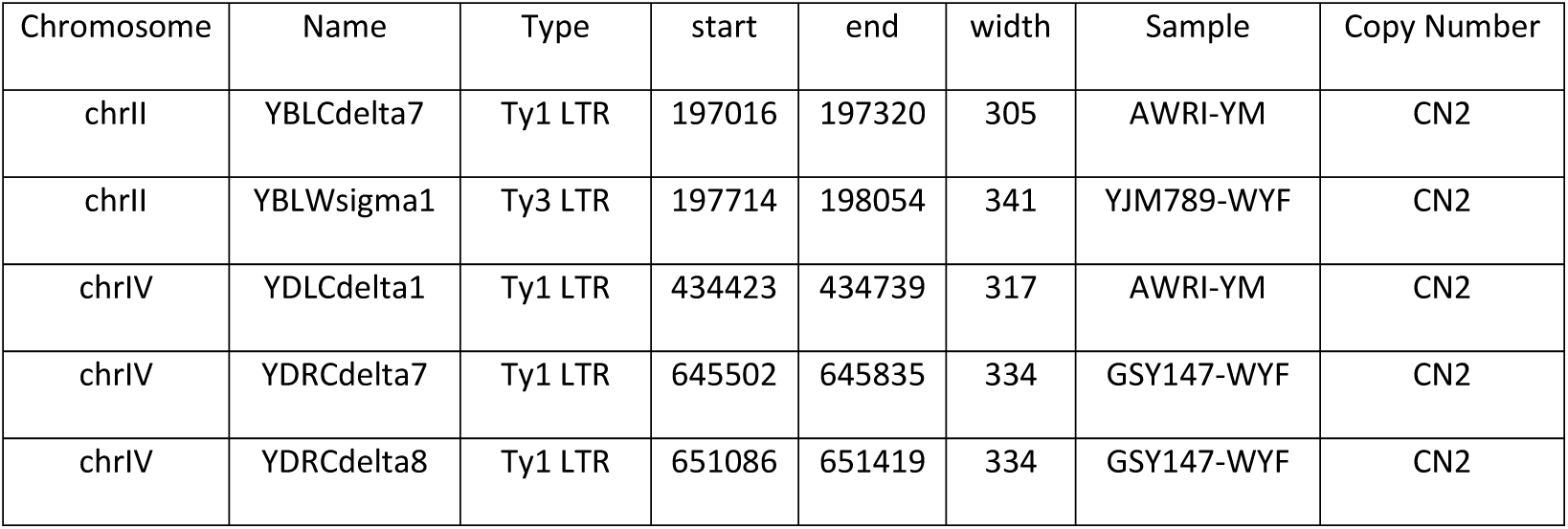
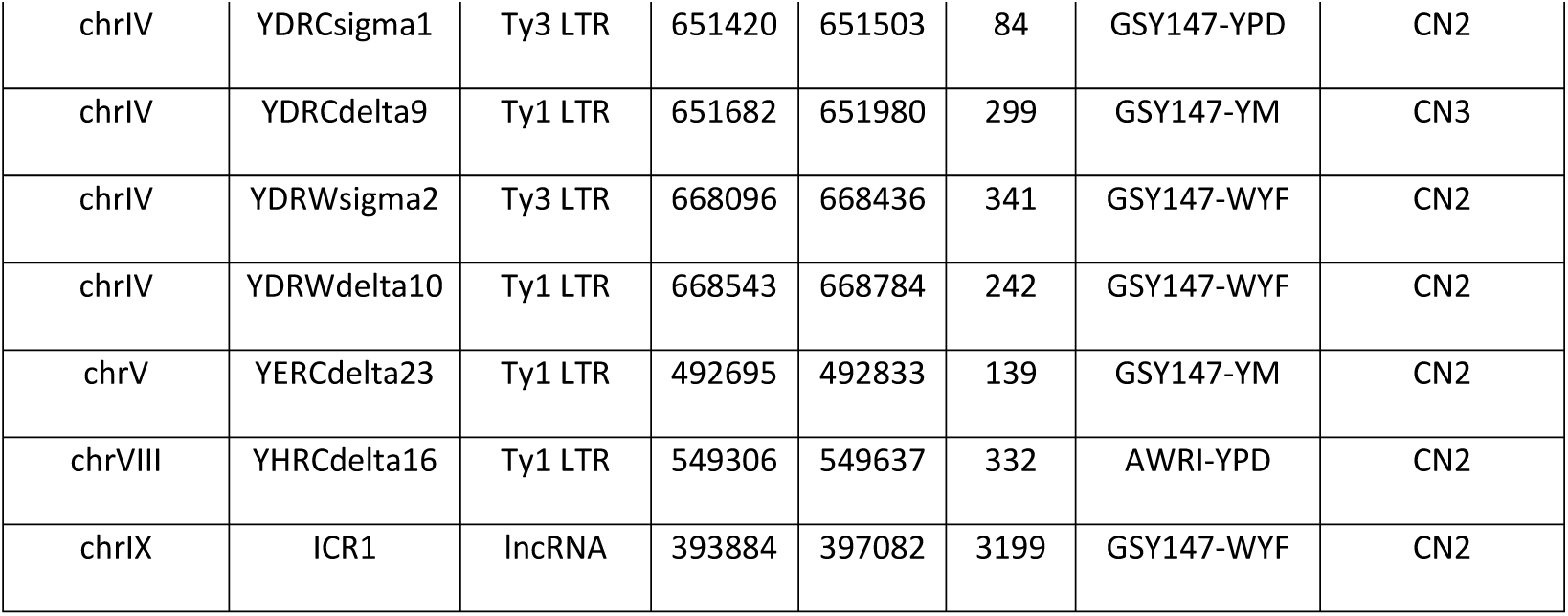
Regions with CNVs flanked by fragile sites in the ILEs

### Transcriptome analysis revealed that glyphosate-tolerance is a polygenic trait

To determine if there was a correlation between the genes that underwent changes in the ILEs and those that are differentially expressed on exposure to Cr41, an RNAseq experiment was carried out. The RNA was extracted and sequenced from cells exposed to Cr41 in minimal media and media supplemented with WYF. The transcriptome analysis was performed on two strains, one of which was sensitive (S288c) to Cr41 exposure and the other resistant (RM11). To decide on which two strains to consider for this study, we also factored in that the two strains had the most variation in the ILE study in terms of number of SNPs and the genes that underwent CNV. S288c, the sensitive strain treated with Cr41, had a much higher number of differentially expressed genes from the RNAseq (Fig 4A). In YM, 1100 genes (S5 Table) and 438 genes (S6 Table) in media supplemented with WYF were differentially expressed in S288c when exposed to Cr41 with a final concentration of 0.25% glyphosate. In RM11, only 58 (S7 Table) and 53 (S8 Table) genes were differentially expressed in YM and media supplemented with WYF, respectively. RM11 differentially expressed few genes in both conditions i.e., minimal media and media supplemented with WYF (Fig 4B) and had a higher correlation with the conditions under which the cells were exposed to Cr41. KEGG pathway enrichment analysis of the differentially expressed genes in S288c revealed that a large number of these genes corresponded to pathways involved in the cell cycle, meiosis, DNA replication, and MAPK signaling pathways, especially in WYF (Fig 4C and 4D). Among many others, *SED1* is one of the cell wall genes that are differentially regulated along with a few transposable elements in S288c, but no SNPs were found in the ILEs. The downregulated genes mainly associated with biosynthesis of secondary metabolites and amino acids in both minimal media and media supplemented with WYF.

**Fig 4.**
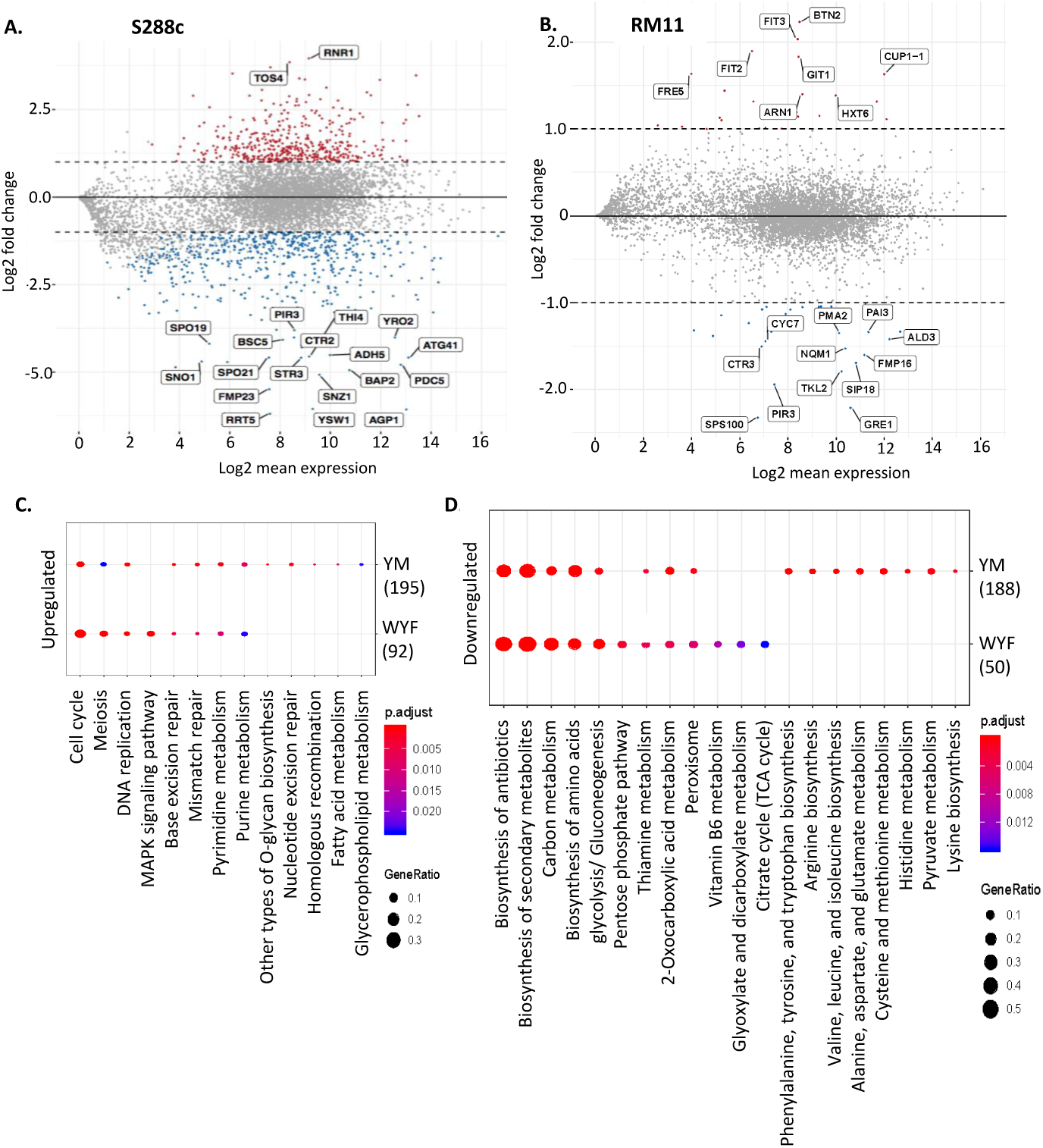
Gene expression analysis of S288c and RM11 on exposure to glyphosate (Credit41) in minimal media A. Dot-plot of significantly up and down-regulated genes in S288c and **B**. in RM11. **C**. KEGG Pathway Enrichment analysis of upregulated genes in S288c, identifying the different pathways involved. **D**. KEGG Pathway Enrichment analysis of downregulated genes in S288c, identifying the different pathways involved.

### S288c cells arrested in G1 on exposure to Cr41 but not pure glyphosate

The RNAseq data showed upregulation of the expression of cell cycle regulator genes encoding proteins that are G1 phase regulators such as Rad53, Cdc28, Nrm1 and Swi4 (Bertoli et al. 2013). Increased expression of these genes suggested that cells were arrested in G1 on exposure to Cr41. To ascertain if it was the glyphosate itself or a cumulative effect of all the additives in CFG that caused cell cycle arrest, cells treated with Cr41 and pure glyphosate were subjected to flow cytometry. RM11 and S288c were grown to log phase and then the asynchronous populations were exposed to 0.25% pure glyphosate and Cr41 (Fig S4). Within 30 minutes of Cr41 exposure, 70% of the population arrested in G1 and remained the same over the course of 6 hours while the culture density was maintained consistently (Fig 5). This was not the case in the untreated and pure-glyphosate treated cells. In these two cases, the populations were evenly distributed across the G1, G2, and S phases. RM11 cells did not show any changes in the cell cycle with respect to the two treatments. It was only the strain that was sensitive to Cr41 exposure, S288c that underwent G1 arrest almost immediately on exposure to Cr41.

**Figure 5.**
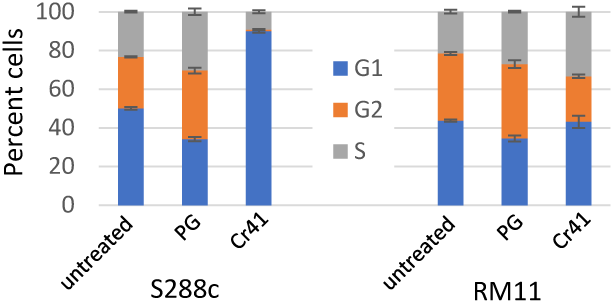
Cell cycle distribution on treatment with glyphosate and commercial glyphosate formulation. Distribution of the cell cycle stages of S288c and RM11 cells on exposure to Credit41 (Cr41) and pure glyphosate (PG) for six hours.

### *sed1Δ* mutants are highly sensitive to Cr41 exposure

The genes commonly affected in the ILE study and differentially expressed in the RNAseq analysis are involved in various pathways, including functions ranging from cell wall proteins, mitochondrial proteins, MAPK related proteins to meiosis (S4 Table). The BY4741 (also an S288c derived strain) knockout collection was used to test the growth phenotype of the different genes as representatives from some of these pathways. Five genes were selected for further characterization, all of which encoded proteins in the cell wall or cell membrane. Det1 plays an integral role in intracellular sterol transport [44]. Pst1 encodes a GPI protein that works with Ecm33 to maintain cell wall integrity [45]. The presence of Emc33 could be the cause for lack of sensitivity in *pst1Δ* mutants on exposure to Cr41. *SED1* is a gene that is expressed as a major stress-induced cell wall glycoprotein [25]. *FLO11* encodes a GPI-anchored cell wall protein, whose transcription is regulated by the MAPK pathway [46]. The *flo11Δ* mutants showed growth defects in minimal media. *VBA5* is a paralog of *VBA3*, and it codes for a plasma membrane protein that plays a role in amino acid uptake [47]. Not all the knockout genes changed growth in response to Cr41 exposure as many of the genes may work in unison to have an overall effect on resistance to the treatment. Another possibility is the specific mutation could have a gain of function effect, which cannot be mimicked by using a knockout collection. One of the cell wall genes that had a definitive response was *SED1*. *SED1* was found in both, the ILE data with its copy number doubled, and expression decreased by 1.475 log2 fold in the transcriptome analysis. The *sed1Δ* mutant was extremely sensitive to Cr41 exposure in both rich and minimal media (Fig 6A). In the S288c ILE strain that developed resistance to Cr41, *SED1* underwent duplication in all media conditions, i.e., minimal media with and without WYF, and rich media. Downregulation of *SED1* gene in YM in S288c (sensitive strain) could contribute to the sensitivity of the cells to Cr41. No mutations were found in *SED1* from the ILE sequencing but there are several insertions and deletions found in RM11, YJM789, and AWRI1631 in reference to S288c (Fig 6B). Sed1 is a stress-induced structural GPI (glycosylphosphatidylinositol) cell wall glycoprotein (Shimoi et al.1998), hence its sensitivity to Cr41 due to the additives could be affecting the integrity of the cell.

**Fig 6:**
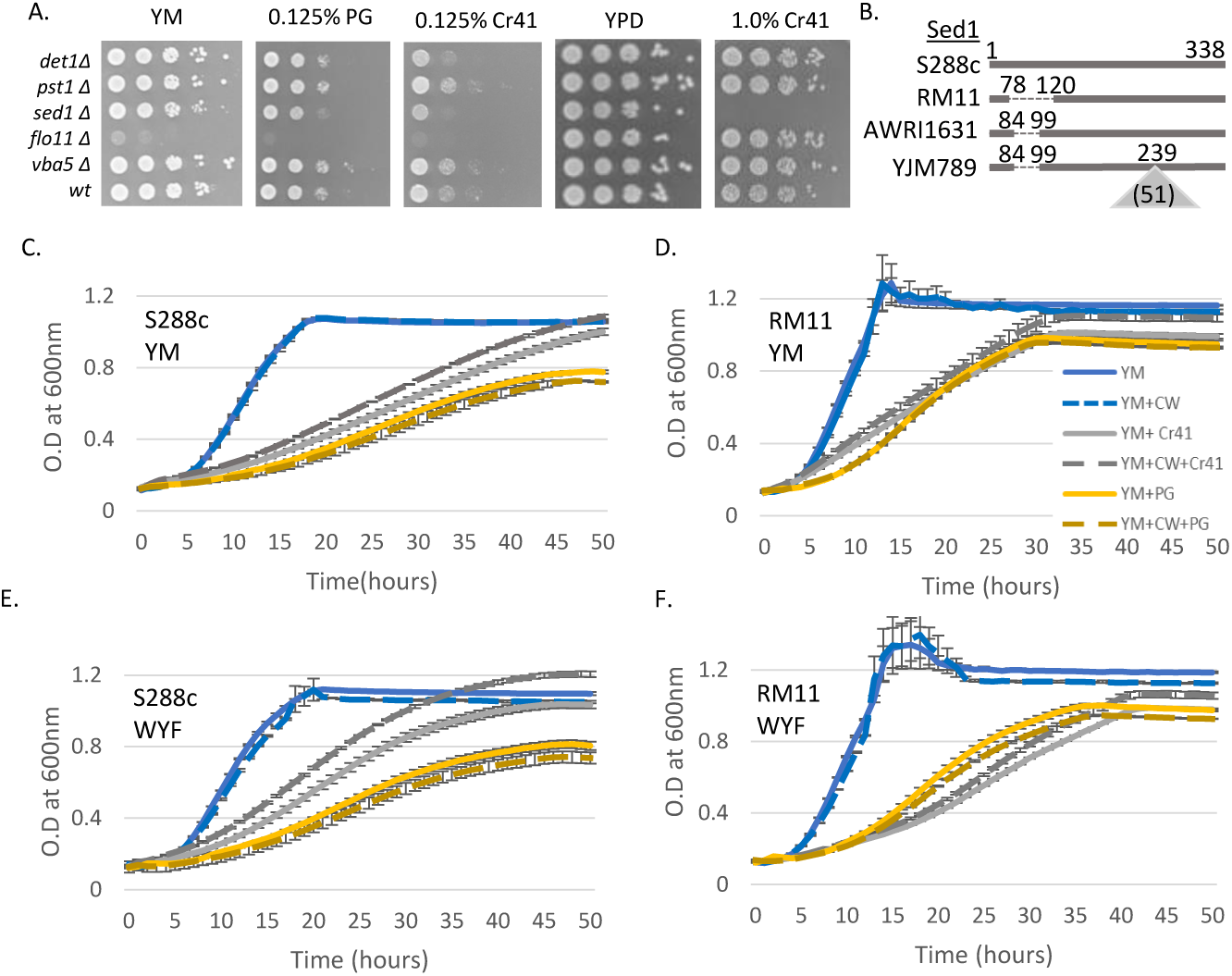
Growth assay on exposure to calcofluor white and glyphosate-based herbicides. **A**. Semi-quantitative growth assay of cell wall mutants. Cells were grown for three days. **B**. Protein alignment of Sed1. Deletions in reference to the S288c are a dashed line and insertions are denoted by a triangle. Amino acid numbers are in reference to S288c with the number of inserted amino acids in parenthesis. Quantitative growth assays of cells grown in minimal media when exposure to calcofluor white (CW), Credit41 (Cr41) and pure glyphosate (PG). OD_600_ was measured for 50 hours in an automatic plate reader. **C**. S288c in YM **D**. RM11 in YM. Yeast were grown in minimal media supplemented with tryptophan (W), tyrosine (Y), phenylalanine (F). **E**. S288c in WYF. **F**. RM11 in WYF.

### Calcofluor white (CFW) rescues growth inhibition of S288c caused by Cr41 exposure

The analysis of affected genes in the evolved strains and the differentially expressed genes gave rise to a significant number of genes that are associated with the cell wall. To test if the cell wall played an important role in the effectiveness of Cr41, one sensitive and one resistant strain that was treated with Cr41 was exposed to Calcofluor white (CFW). CFW is known for its property of inducing cell wall stress, as it is a chitin antagonist and results in increased deposition of chitin making the cell walls thicker (Liesche, et al. 2015; Roncero and Duran 1985). A quantitative liquid growth assay was carried out using S288c and RM11. The growth of RM11 was inhibited by pure glyphosate and to a greater extent by Cr41. Treating the RM11 samples with CFW did not have a significant effect in YM for Cr41 or PG (p value=0.19 and 0.503). In WYF, it was significant with Cr41 (p-value=0.011) but not PG (p-value=0.08). All strains closely related to S288c showed a higher sensitivity to pure glyphosate compared to Cr41 over the first 50 hours (Fig S5). Treating S288c cells in YM exposed to pure glyphosate with CFW did not have a significant effect (p-value = 0.378) (Fig 6C-E). However, CFW significantly alleviated growth inhibition caused by Cr41 (p-value = 0.024). Cells treated with CFW have increased cell wall volume by about 30% and the wall/ cell ratio also increases significantly (Liesche et al. 2015). The increase in cell wall thickness could be the main contributor to the rescue of cells treated with Cr41. This indicated that the additives in Cr41 along with the glyphosate have a cumulative effect on the cell wall of sensitive strains such as S288c, which was alleviated by CFW treatment. The increase in cell wall thickness on CFW treatment helps the cells withstand the effects of Cr41, also it may reduce the amount of Cr41 entering into the cell. However, treating cells exposed to only pure glyphosate with CFW does not result in alleviation of growth inhibition. This implies that CFW is not rescuing cells from the glyphosate in Cr41 but the alleviating the effects of the additives on the cell wall.

## Discussion

Glyphosate-based herbicides are most commonly used around the world because of the specificity of glyphosate, in acting solely on the aromatic amino acid pathway, that is absent in humans and other eukaryotes. The additives and surfactants present in the commercial formulations are chosen due to their intrinsic inert and non-toxic properties. There are variations from one CFG to the other in terms of these additives which results in different extents of the effectiveness of the herbicide [13]. The supposed non-activeingredients that are added in CFG, enhance the potency of glyphosate and were not inert as seen by changes in the transcriptome. A crucial part is the study of all the changes occurring within the cell to make the herbicide more effective due to the presence of additives and surfactants. The additives in the CFG along with other functions play a key role in the entry of the active ingredient into the cell. This is done by first compromising the first protective physical barrier that they come in contact with, the cell wall. The effectiveness varies based on the structure and composition of the cell wall along with the alleles of the different mannoproteins that the cell contains. When the concentrations and combinations of additives change, this also changes their cumulative effect on the cell. When multiple CFG were tested on different strains of yeast, this resulted in variation in their growth phenotype. The strains chosen for this study have been isolated from different environments as prior exposure was suspected to have an effect of the extent of tolerance to CFG. The agricultural isolates (RM11, AWRI1631) were more resistant to CFGs and glyphosate compared to the laboratory strain and the clinical isolate (YJM789). The agricultural isolates may have come in contact with glyphosate or similar herbicides in the past, which could have led to the development of resistance mechanisms based on past exposure. The lab strain and clinical isolate are not in environments commonly exposed to CFG, making it less likely for them to have been exposed in the past, contributing to their sensitivity.

The genetic variation between these strains was used to gain a better understanding of the effects of the CFGs as a whole at the genome and transcriptome level. In case of some stressors, adaptation can occur through a specific route depending on the regions of the genome most effected by the stressor. Response to CFG is a polygenic trait as the different components of the herbicide act together to affect many genes that contribute to various pathways. Hence adaptation can result from small modifications that occur in many different genes as a cumulative effect, as well as one particular change that could have caused a population bottleneck and dominated the population. Large sections of the genome were found to have undergone CNV, many of which were flanked by Ty elements and ncRNAs. Regulatory elements have been shown to play an important role in combatting stressors that the cell encounters [50]. *S*. *cerevisiae* has five families of LTR retrotransposons, and they comprise about 3% of the entire genome [51]. Mutations in certain DNA repair genes results in expression of common fragile sites, which when induced often undergo translocations, deletions, duplications, etc. [52]. The ILE strains contained mutations in genes involved in many pathways, some of which were contributors to DNA repair mechanisms (such as *RAD53, UME6*, and *MSH3*) as well. Any of these alterations could have induced fragile site expression, in turn resulting in duplication of genes present between two Ty elements. Having the Ty elements in these regions provided the opportunity for these genes to undergo duplication, and those with a beneficial effect were retained as the population progressed. Over the course of serial passaging of these strains, the duplicated regions along with the effects of SNPs in some of the genes may have provided the advantage that the cells needed to overcome sensitivity to Cr41. To assess if there was an overarching contribution by a particular gene that underwent modification or it was the cumulative effect of many of the changes, some of the genes were tested using the knockout collection. This led to the observation that Sed1 is an important contributor to the cell’s resistance of CFG.

The most drastic effects of glyphosate-based herbicides on the cell apart from the aromatic amino acid pathway was on the cell wall. The cell wall is the mode of entry into the cell and a vital structure in maintaining the osmotic integrity of the cell. Changing the combinations or concentrations of additives in an herbicide alters the mode of entry of the active component into the cell. In our previous study, we have shown that proteins such as Dip5, the aspartic and glutamic acid permease [10] are involved in the import of some glyphosate into the cell. Dip5 was found by comparing S288c and YJM789 genomes with QTL analysis. Therefore only genes that had genetic variation between these strains would be detected and there could be permeases/ transports that also regulate glyphosate that are genetically identical between strains that would not be identified in QTL analysis. The effect of pleiotropic drug response genes is more evident in rich media. *PDR5* is the most polymorphic gene in yeast [53]. Pdr5 shares 96% amino acid similarity between S288c and YJM789. Whereas Pdr5 is 99.7% similar between S288c and RM11, which may have led to the masking of its contribution to this study. Sed1 is a GPI-cell wall protein that is highly expressed when the cells are in stationary phase and is a required protein for cells in this phase when they are under stress (Shimoi et al. 1998). *SED1* is not very highly conserved across strains and a deletion leading to a similarity of only 87.87% between the S288c and RM11 alleles. Like Pdr5, the role of Sed1 appears to be primarily aimed at dealing with the effects of the additives in Cr41. Like the *pdr5* mutant, the *sed1* was very sensitive on YPD which has higher levels of Cr41 to induce growth inhibition. The MAPK pathway along with Hog1 genes coregulate genes involved in maintaining the integrity of the cell and cell wall. Several Hog1 MAP kinase genes (*FUS3, STE5, BMH2*, and *AFR1*) contained gene duplications in the evolved strains. This pathway is activated under conditions of hyperosmotic stress and is usually accompanied by differential expression of various GPI-cell wall proteins. The CFG as a whole could be inducing hyperosmotic stress resulting in the activation of the Hog1 MAP kinase pathway which could also contribute to arresting in G1 phase (Escoté, et al. 2004).

In this study, we highlight the importance of studying different CFG to gain a better understanding of all the pathways affected on exposure to Cr41 and to recognize the various non-target adaptation mechanisms. Humans have proteins that have structure and functional similarity to those found in yeast. As this herbicide is used so extensively on produce that is used for human consumption, it is important that we understand the effects of the chemicals we are being exposed to. Recent studies have shown the presence of glyphosate and its metabolites in urine samples in humans [55,56] through the levels of glyphosate humans are exposed to through food consumption is much lower than those used in this study to treat yeast cells. Without analyzing the effects at the molecular level, it is difficult to predict if there will be any long-term effects. Not seeing any effects immediately is not necessarily an indication that there would not be any effects after long-term exposure. Hence, it is crucial to study the effects of different CFG to be able to regulate the use of additives and surfactants.

## Materials and Methods

### Variations in growth phenotype

A semi-quantitative growth assay was carried out on exposure of AWRI1631, RM11, YJM789, and S288c, to different formulations of glyphosate-based herbicides. The media used for the study was nutrient-rich (YPD) and nutrient minimal (YM) solid media. The rich media is comprised of yeast extract, peptone, and dextrose; as for the minimal media, it consists of yeast-nitrogen base and 2% dextrose. The minimal media was supplemented with aromatic amino acids, namely 20 µg/ml tryptophan (W), 30 µg/ml tyrosine (Y) and 50 µg/ml phenylalanine (F). It was also supplemented with 100 µg/ml aspartic (D) acid for certain conditions. This assay was performed as previously described in [10], as follows. These were exposed to the glyphosate-based herbicides and the concentration of glyphosate was standardized to 1.0% in rich and 0.15% in minimal media respectively. The different formulations used were, Compare and Save (CAS), WeedPro (WP), Super Cncentrated (SC), Credit 41 (Cr41) and pure glyphosate as a control.

Quantitative growth analysis was carried out using a TECAN M200, automatic plate reader [57]. The growth of cells in liquid media was measured every 1 hour at 600nm under shaking conditions. Along with Credit41 and pure glyphosate, the cells were treated with 5mM calcofluor white, to measure the progression of their growth over 50 hours.

### Gene expression analysis

RNA was extracted from S288c and RM11 cells grown in minimal media with and without supplementing WYF. The cells were treated with 0.25% Credit41 for 90 minutes with 5 replicates of each condition. The samples were then washed, and the total RNA was extracted using hot phenol method [58]. Paired-end cDNA libraries were built using the RNA extracted using Epicenre’s ScriptSeq Yeast kit. The sequencing was performed using 76bp paired-end reads on the Illumina HiSeq platform resulting in 5-8.5 million read pairs.

Salmon v0.13.1 was used to estimate the transcript level abundance from the RNA seq-read data [59]. To do so, indexing of the S288c reference transcriptome was first carried out. This index was used by Salmon against each sample to generate quant.sf files containing the length, abundance in terms of Transcripts Per Million (TPM) and the estimated number of reads for each transcript(https://www.ncbi.nlm.nih.gov/geo/query/acc.cgi?acc=GSE135473 secure token: arkxmokwdlufjir). The differential expression of genes between the two strains and each condition was calculated using DESeq2 v1.8.1. DESeq2 was also used to generate a sample to sample distance map (Fig S3). The program clusterProfiler v3.10.0 was used to carry out KEGG Pathway Enrichment Analysis, to identify the pathways involved, based on the clustering of genes.

### Flow cytometry

To measure the cell cycle arrest on Credit41 and pure glyphosate exposure, S288c and RM11 cells were subjected to flow cytometry after exposure. The cultures were started and allowed to grow to mid-log phase from an overnight culture. Once the cells reached mid-log phase, they were treated with 0.25% Credit41 and pure glyphosate, in triplicate with and without supplementing WYF in minimal media. Cells were collected at multiple time points (0 mins, 30 mins, 90 mins, 3 hrs, and 6 hrs) over the next 6 hours. The cells were maintained in log-phase throughout the collections, by replenishing with fresh media as needed. The cells were harvested and fixed in 70% ethanol for 2 days [60]. The cells were then washed and treated with RNAse solution for 8-12 hours at 37°C. The RNAse solution comprised of a mixture of 2mg/ml RNAse A, 50mM Tris pH 8.0, and 15mM sodium chloride that had been boiled for 15 minutes and cooled to room temperature. The cells were then treated with protease solution (5 mg/ml pepsin and 4.5 µl/ml concentrated HCl) for 20 minutes at 37°C. The cells were then stored in 50 mM Tris pH 7.5 at 4°C, until the day of the analysis. Right before flow cytometry analysis, the cells were sonicated at low levels to separate cells. 50ul of the sonicated mixture was transferred into 1ml of 1uM Sytox Green in 50ml Tris pH 7.5. The cells were analyzed on an LSRFortessa, using the FITC channel. The results were analyzed, and the percentage of cells in each stage of the cell cycle was estimated using FCS Express 6.0. The analysis was carried out using the multi-cycle DNA histogram and ‘+S order=1 model’ was selected based on the lowest Chi-square value.

### Whole-genome sequencing of In-Lab evolutions (ILEs)

In-lab evolutions were carried out by exposing cells to Credit41 over a long period of time until a resistant population was isolated. Two sensitive (S288c and YJM789), and two resistant (RM11 and AWRI1631) strains were used for this study. The cells were evolved in minimal media with 0.25% Cr41, with and without supplementing WYF. The cells evolved in rich media were treated with 1.0% Cr41. A control group was evolved in media in the absence of Credit41 to account for mutations that occur due to the procedure, rather than the Credit41 treatment. A single colony was used to start a saturated overnight culture, that was then used as a starter culture to inoculate the different conditions in triplicate. Each passage was grown 2-3 days in media till saturation. After which 1% of the culture was transferred into fresh media. The cells were subjected to 6 passages and tested for resistant populations by plating 10-fold dilutions on solid media. The resistant populations were streaked on plates to isolate single colonies. The resistant colonies were selected based on size and shape. They were then passaged for 2 passages in media lacking Credit41, to ensure the resistance did not rely solely on epigenetic mechanisms. The genomic DNA was then extracted using the 96 well genomic DNA extraction kit, and the phenol-chloroform extraction technique was also used [61].

The whole-genome sequencing data for all 4 strains were aligned to S288c reference sequence (release R64-2-1) by creating an index and using gatk-4.1.1.0 to carry out the alignment. Samtools was used to convert the SAM files to BAM files, from which duplicates were removed. Using gatk HaplotypeCaller, variant calling was carried out to identify all the SNPs and generate vcf files. Using the generated vcf files, PCA variants were identified using pcadapt as described in (Hugh et al. 2019) (https://data.mendeley.com/datasets/ts5syfw38r/draft?a=e2170360-459e-4a8c-b1b3-d89f4b530fdf) (doi 10.17632/ts5syfw38r).

## Acknowledgments

We thank Angela Lee for the yeast knockout collection. We would like to thank the WVU Flow Cytometry Core (Fortessa S10 grant: OD016165) and the Genomics Core Facility for their services. This work was supported by NSF MCB-1614573 and WVU Senate Research Grant program. R-14-003.

## Author Contributions

JEGG conceived and directed this study. AR carried out all the experiments and wrote the manuscript. AR and AP mapped, called CNVs and SNPs, and quantitated the transcriptomic and genomic sequences. The authors declare that they have no competing financial interests.

---

**S1 Fig:**
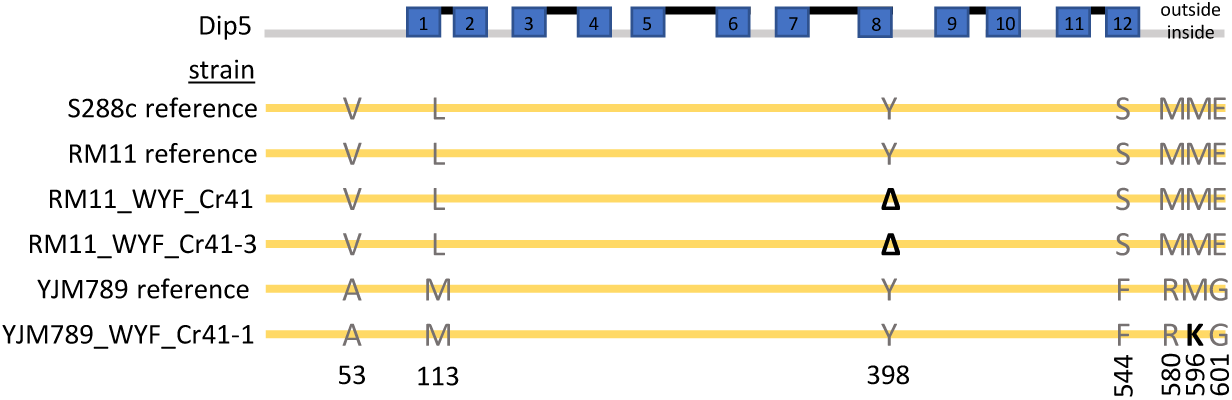
Alignment of ILE strains containing mutations in *DIP5*. Three of the ILE strains contained mutations in the *DIP5* gene. These were aligned to highlight all the polymorphisms in grey and mutations in bold with reference to the S288c and to each respective reference genome. The protein topology diagramed at the top is represented by black lines as the extracellular sequence and grey is the intercellular sequence. The blue boxes were predicted by TMHMM to be transmembrane domains (TM). The sequences for each TM of Dip5^S288c^ are as follows TM1 91-113, TM2 120-142, TM3 157-179, TM4 200-219, TM5 234-253, TM6 286-308, TM7 323-345, TM8 376-398, TM9 423-445, TM10 455-477, TM11 500-522, and TM12 532-551.

**S2 Fig:**
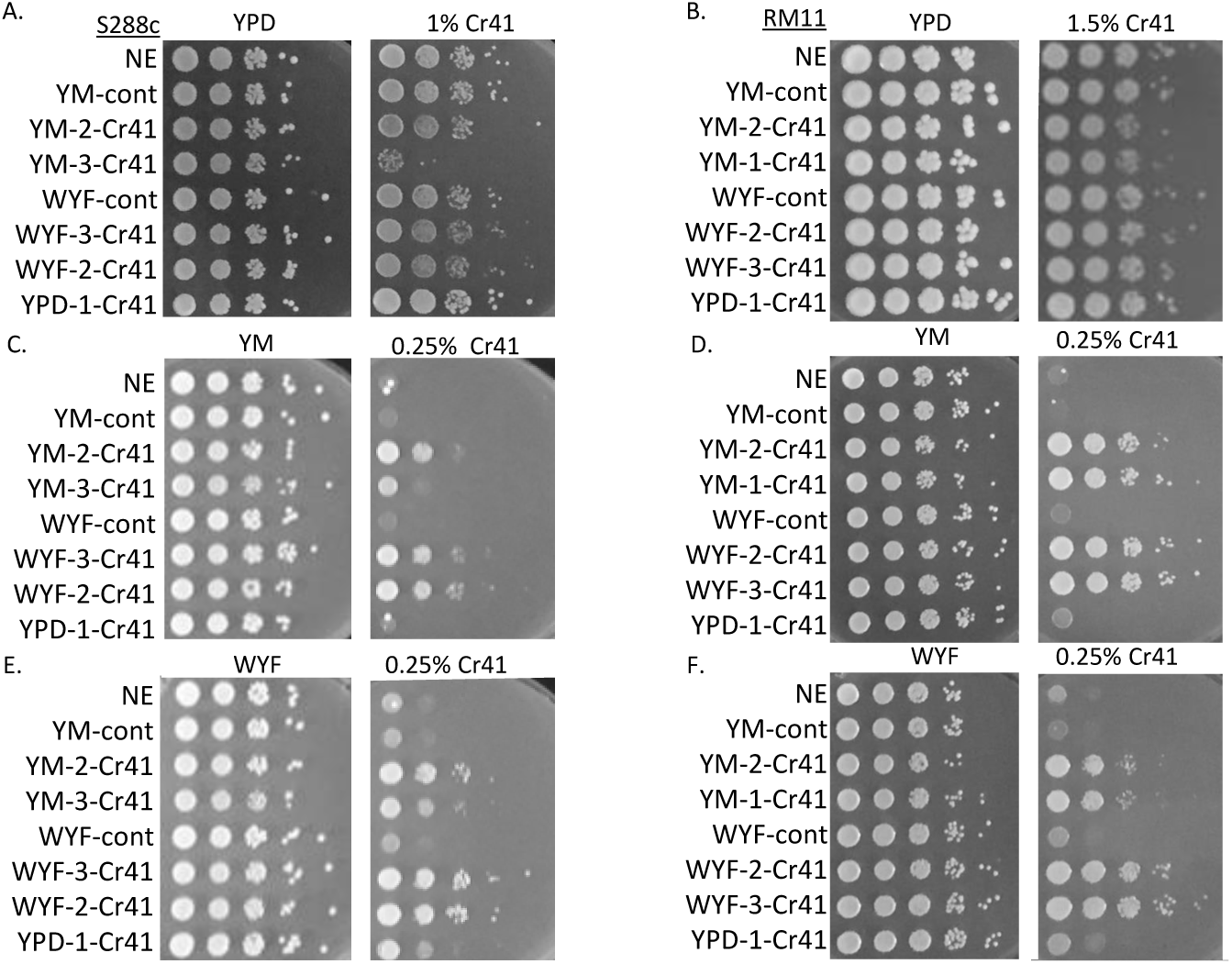
Cross-resistance was conferred in strains evolved in minimal media and WYF. **A**. Strains evolved under different conditions were tested for resistance in rich media with Cr41 for S288c and **B**. RM11. **C**. Strains evolved under different conditions were tested for resistance in minimal media with Cr41 in S288c and **D**. RM11. **E**. Strains evolved under different conditions were tested for resistance in minimal media supplemented with WYF and treated with Cr41 in S288c and **F**. RM11.

**S3 Fig:**
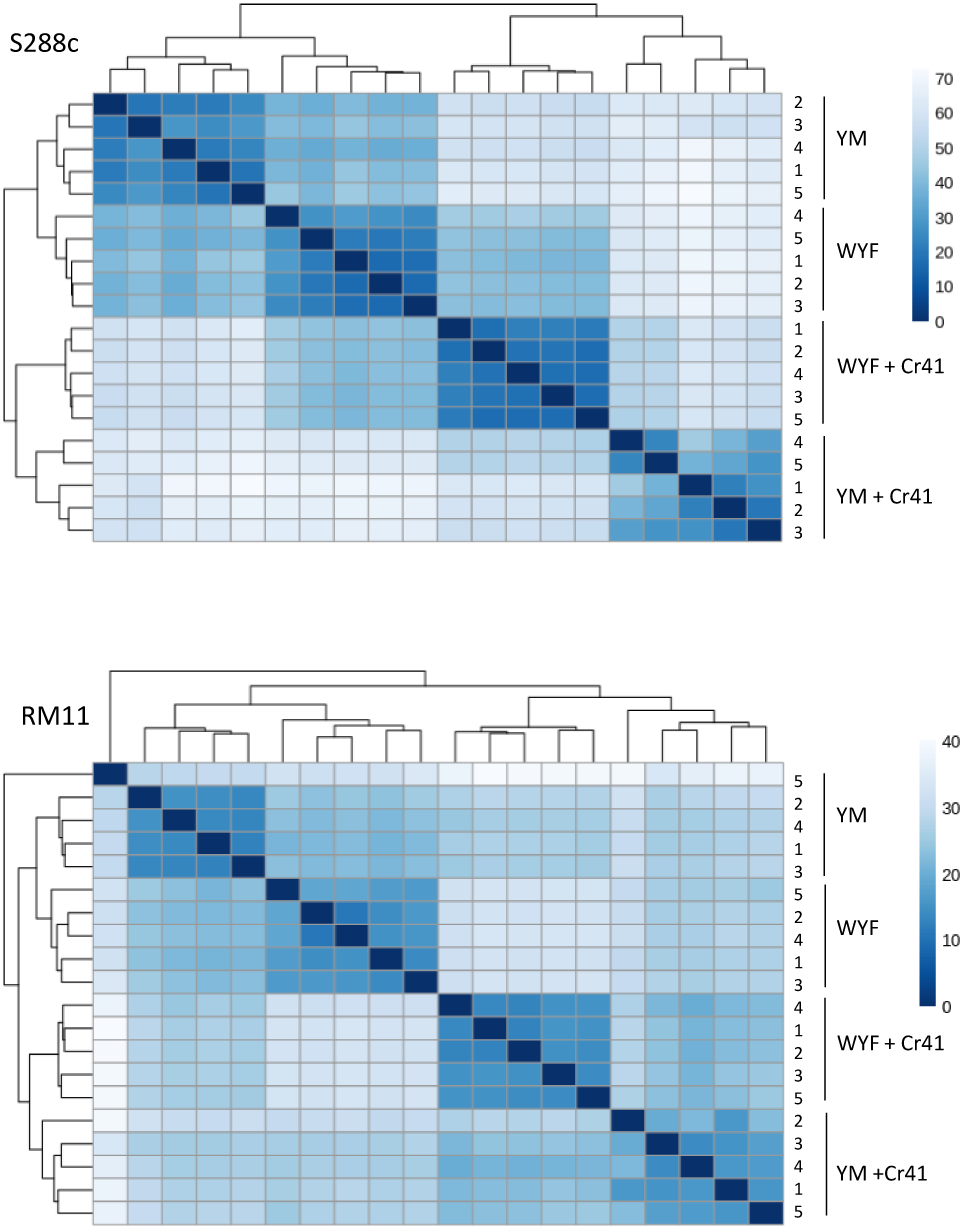
Sample to sample distance estimation using DESeq2. The distance map shows clustering of biological replicates of RNAseq samples for **A**. S288c and **B**. RM11.

**S4 Fig:**
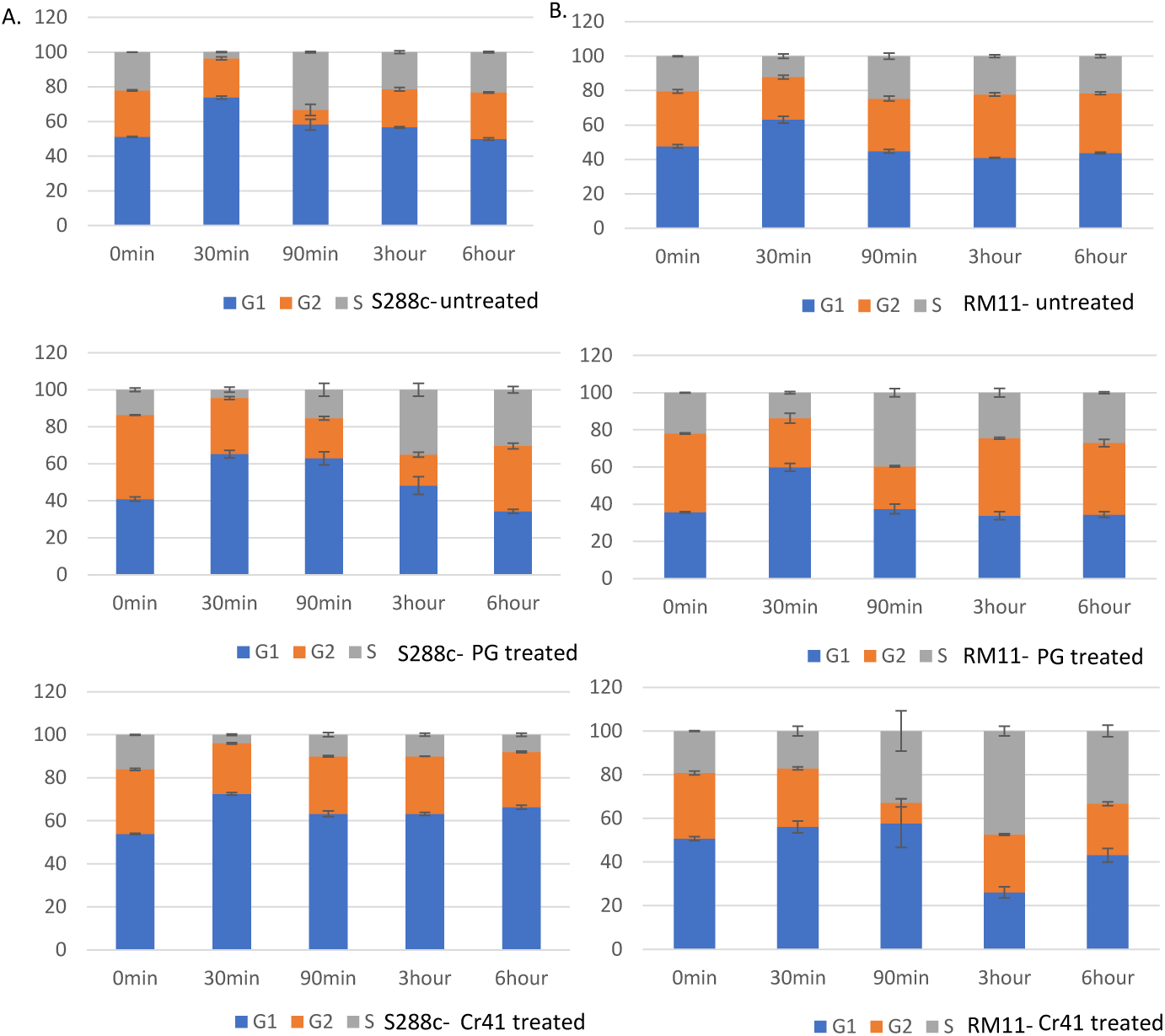
Flow cytometry of S288c strains show arrest in the G1 phase of the cell cycle on treatment with Cr41. **A**. Cell cycle distribution of S288c shows arrest in case of treatment with Cr41 but no arrest when treated with pure glyphosate. **B**. Cell cycle distribution of RM11 shows no effect on treatment with Cr41.

**S5 Fig:**
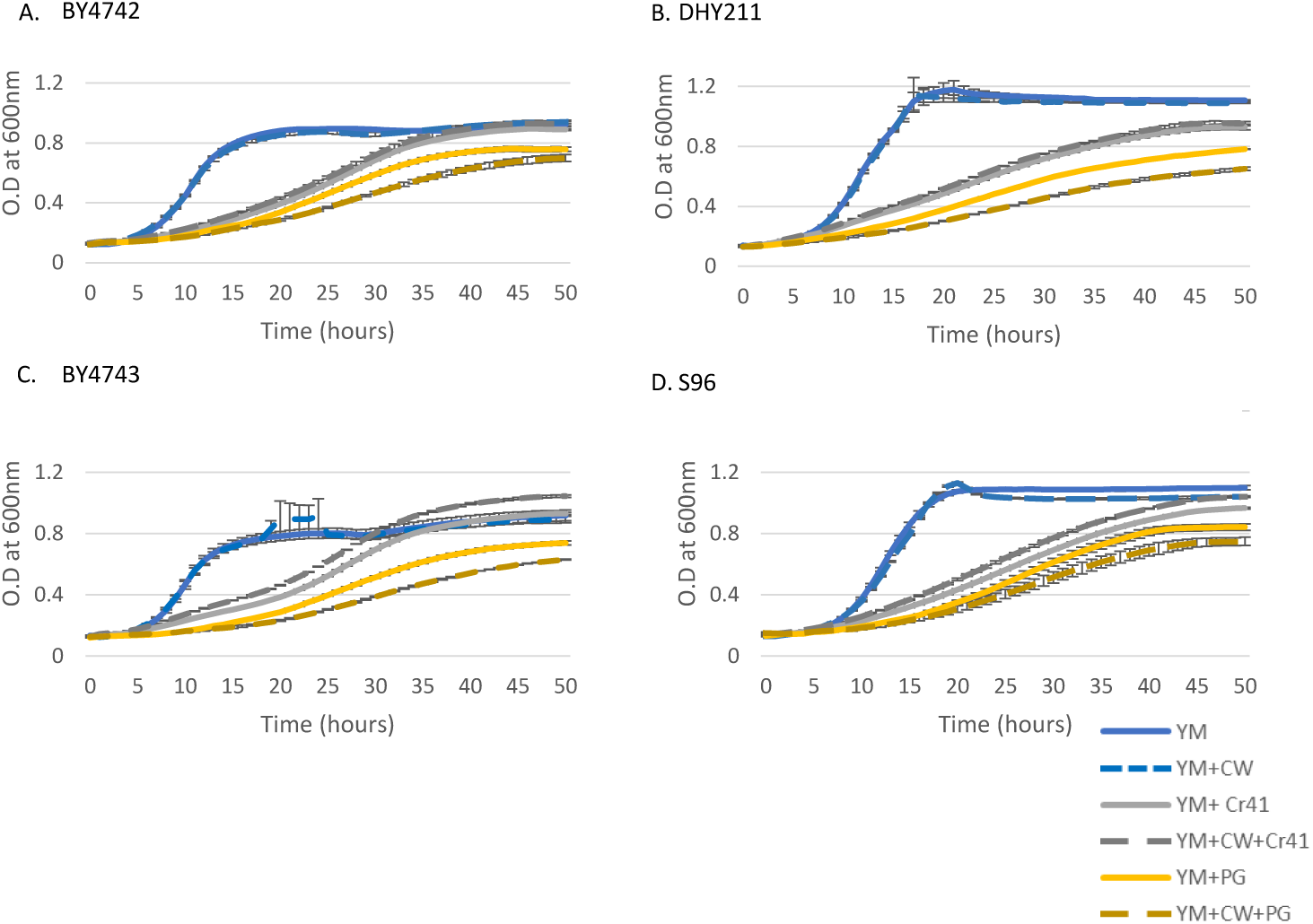
Pure glyphosate inhibits growth to a greater extent than Cr41 only in strains closely related to S288c. S288c related cells were grown in minimal media, their growth was measured on exposure to calcofluor white (CW), Credit41 (Cr41) and pure glyphosate (PG) **A**. BY4742 **B**. DHY211 **C**. BY4743 **D**. S96

